# RGS14 restricts plasticity in hippocampal CA2 by limiting postsynaptic calcium signaling

**DOI:** 10.1101/297499

**Authors:** Paul R. Evans, Paula Parra-Bueno, Michael S. Smirnov, Daniel J. Lustberg, Serena M. Dudek, John R. Hepler, Ryohei Yasuda

**Affiliations:** Max Planck Florida Institute for Neuroscience, Jupiter, FL, 33458, USA; Department of Pharmacology, Emory University School of Medicine, Atlanta, GA 30322, USA; Neurobiology Laboratory, National Institute of Environmental Health Sciences, National Institutes of Health, Research Triangle Park, NC, 27709, USA

## Abstract

Pyramidal neurons in hippocampal area CA2 are distinct from neighboring CA1 in that they resist synaptic long-term potentiation (LTP) at CA3 Schaffer Collateral synapses. Regulator of G Protein Signaling 14 (RGS14) is a complex scaffolding protein enriched in CA2 dendritic spines that naturally blocks CA2 synaptic plasticity and hippocampus-dependent learning, but the cellular mechanisms by which RGS14 gates LTP are largely unexplored. A previous study has attributed the lack of plasticity to higher rates of calcium (Ca^2+^) buffering and extrusion in CA2 spines. Additionally, a recent proteomics study revealed that RGS14 interacts with two key Ca^2+^-activated proteins in CA2 neurons: calcium/calmodulin, and CaMKII. Here, we investigate whether RGS14 regulates Ca^2+^ signaling in its host CA2 neurons. We find the nascent LTP of CA2 synapses due to genetic knockout (KO) of RGS14 in mice requires Ca^2+^-dependent postsynaptic signaling through NMDA receptors, CaMK, and PKA, revealing similar mechanisms to those in CA1. We report RGS14 negatively regulates the long-term structural plasticity of dendritic spines of CA2 neurons. We further show that wild-type (WT) CA2 neurons display significantly attenuated spine Ca^2+^ transients during structural plasticity induction compared with the Ca^2+^ transients from CA2 spines of RGS14 KO mice and CA1 controls. Finally, we demonstrate that acute overexpression of RGS14 is sufficient to block spine plasticity, and elevating extracellular Ca^2+^ levels restores plasticity to RGS14-expressing neurons. Together, these results demonstrate for the first time that RGS14 regulates plasticity in hippocampal area CA2 by restricting Ca^2+^ elevations in CA2 spines and downstream signaling pathways.

**Significance Statement:** Recent studies of hippocampal area CA2 have provided strong evidence in support of a clear role for this apparently plasticity-resistant subregion of the hippocampus in social, spatial, and temporal aspects of memory. Regulator of G Protein Signaling 14 (RGS14) is a critical factor that inhibits synaptic plasticity in CA2, but the molecular mechanisms by which RGS14 limits LTP remained unknown. Here we provide new evidence that RGS14 restricts spine calcium (Ca^2+^) in CA2 neurons and that key downstream Ca^2+^-activated signaling pathways are required for CA2 plasticity in mice lacking RGS14. These results define a previously unrecognized role for RGS14 as a natural inhibitor of postsynaptic Ca^2+^ signaling in hippocampal area CA2.

## Introduction

Pyramidal neurons in hippocampal area CA2 differ markedly from neighboring subfields in that synaptic long-term potentiation (LTP) is not readily induced (Zhao et al., 2007). A number of genes are selectively expressed in CA2 pyramidal neurons (Lein et al., 2005, 2007; Cembrowski et al., 2016), and a few of these proteins have been shown to contribute to the atypical plasticity features of CA2 (Simons et al., 2009, 2012; Lee et al., 2010; Caruana et al., 2012; Pagani et al., 2015; Carstens et al., 2016; Dudek et al., 2016). Regulator of G Protein Signaling 14 (RGS14) is one such gene that is highly expressed in CA2 pyramidal neurons (Evans et al., 2014), where it is enriched postsynaptically in dendrites and spines (Lee et al., 2010; Squires et al., 2017). RGS14 knockout (KO) mice possess an unusually robust capacity for LTP in CA2, which is absent in wild-type (WT) mice, and exhibit markedly enhanced spatial learning (Lee et al., 2010). Thus, RGS14 acts as a critical factor restricting synaptic plasticity in CA2 pyramidal neurons and hippocampal-based learning and memory.

RGS14 is a complex scaffolding protein with an unconventional protein architecture that allows it to integrate G protein signaling and ERK/MAPK signaling (Shu et al., 2010; Vellano et al., 2013; Evans et al., 2015). RGS14 interacts with active Gαi/o-GTP subunits through an RGS domain and inactive Gαi1/3-GDP subunits through its GPR motif (Cho et al., 2000; Traver et al., 2000; Hollinger et al., 2001). RGS14 also binds active HRas-GTP and Raf kinases through the tandem Ras-binding domains (Kiel et al., 2005; Shu et al., 2010; Vellano et al., 2013). The nascent plasticity of CA2 synapses in RGS14 KO mice requires ERK signaling (Lee et al., 2010), but additional cellular mechanisms have yet to be explored.

An independent study has attributed the lack of plasticity in CA2 pyramidal neurons to smaller spine Ca^2+^ transients due to higher Ca^2+^ extrusion rate and buffering compared to CA1 and CA3 pyramidal neurons (Simons et al., 2009), and Ca^2+^-dependent mechanisms underlie some forms of CA2 synaptic potentiation (Simons et al., 2009; Pagani et al., 2015). Elevations in postsynaptic Ca^2+^ are critical to induce activity-dependent LTP in CA1 hippocampal neurons (Lynch et al., 1983; Malenka et al., 1988) as well as in CA2 (Simons et al., 2009). Ca^2+^ binds calmodulin (Ca^2+^/CaM), which then potently activate many plasticity-inducing pathways, including CaMK, Ras/ERK, and PKA (Malenka et al., 1989; Abel et al., 1997; Otmakhova et al., 2000; Yasuda et al., 2006; Lee et al., 2010; Tang and Yasuda, 2017). A very recent proteomic analysis found that RGS14 natively interacts with proteins that regulate actin binding, CaMK activity, and CaM binding, in mouse brain (Evans et al., 2018). From these candidate interactors, two new *bona fide* RGS14 binding partners were identified that are Ca^2+^-activated and required for plasticity: Ca^2+^/CaM and CaMKII. Moreover, proximity ligation assays revealed that RGS14 interacts with CaM and CaMKII in murine hippocampal CA2 neurons (Evans et al., 2018). However, to date, no evidence has functionally linked RGS14 to Ca^2+^ signaling pathways required for LTP induction. Therefore, we investigated whether RGS14 modulates key Ca^2+^-stimulated pathways to restrict plasticity in CA2 hippocampal neurons.

Here, we performed electrophysiology and two-photon fluorescence imaging experiments in brain slices from RGS14 WT and KO littermates. Using a reporter mouse line to localize CA2 dendrites to perform field recordings, we found the nascent LTP of CA2 synapses in adult RGS14 KO mice requires Ca^2+^-dependent signaling through NMDARs, CaMK, and PKA. Given the similarity to mechanisms supporting structural plasticity of dendritic spines in CA1, we next discovered that RGS14 limits CA2 spine structural plasticity. We found that RGS14 likely accomplished this by attenuating spine Ca^2+^ levels during spine plasticity induction as spines of CA2 neurons from WT mice showed smaller Ca^2+^ transients during synaptic stimulation compared to CA2 neurons from RGS14 KO mice. Lastly, we found that acute overexpression of RGS14 in brain slices from RGS14 KO mice can “rescue” the impairment in long-term spine plasticity; sustained spine plasticity can be restored to RGS14-expressing neurons by elevating extracellular calcium levels suggesting that Ca^2+^ signaling is central to this process. Our results presented here define a new role for RGS14 as a novel regulator of postsynaptic Ca^2+^ signaling and identify a new mechanism upstream of the Ras-ERK pathway whereby RGS14 exerts control over plasticity signaling in CA2 pyramidal neurons.

## Materials and Methods

### Animals

Animals in all experiments were house under a 12h:12h light/dark cycle with access to food and water *ad libitum*. All experimental procedures conform to US NIH guidelines and were approved by the animal care and use committees of the Max Planck Florida Institute for Neuroscience and the National Institute of Environmental Health Sciences. RGS14 KO mice were generated and maintained as previously described (Lee et al., 2010). Both male and female RGS14 WT/KO animals were used in all experiments. Reporter mice expressing enhanced green fluorescent protein (EGFP) in CA2 pyramidal neurons under the Amigo2 promoter (Amigo2-EGFP; Tg(Amigo2-EGFP)LW244Gsat; RRID:MMRRC_033018-UCD) were crossed with RGS14 WT/KO mice to label CA2 dendrites for field recordings.

### Acute slice preparation

Adult RGS14 WT or KO;Amigo2-EGFP+ mice (P20-P50) were sedated by isoflurane inhalation, and perfused intracardially with a chilled choline chloride solution. Brains were removed and placed in the same choline chloride solution composed of 124 mM choline chloride, 2.5 mM KCl, 26 mM NaHCO_3_, 3.3 mM MgCl_2_, 1.2 mM NaH_2_PO_4_, 10 mM glucose and 0.5 mM CaCl_2_, pH 7.4 equilibrated with 95% O_2_/5% CO_2_. Coronal slices (400 µm) were prepared on a vibratome (Leica VT1200), and slices were maintained in a submerged holding chamber at 32°C for 1h and then at room temperature in oxygenated ACSF.

### Extracellular Recordings and LTP protocol

Experiments were performed at room temperature (~21°C), and submerged slices were perfused with oxygenated ACSF containing 2 mM CaCl_2_, 2 mM MgCl_2_ and 100 µM picrotoxin. One or two glass recording electrodes (resistance ~4 MΩ) containing the same ACSF solution was placed in the stratum radiatum of CA2 or CA1 respectively (~100–200 µm away from the soma) while stimulating Schaffer Collateral fibers with square current pulses (0.1 ms duration) using a concentric bipolar stimulation electrode (FHC) every 30 s. For clarity of display, every other data point was removed from averaged time course field recording experiments; all data points were included in the statistical analyses. Area CA2 was detected by Amigo2-EGFP fluorescence. The initial slope of the EPSP was monitored with custom software. The stimulation strength was set to ~50% saturation. LTP was assessed by applying three sets of high frequency stimuli (100 Hz, 1 s) with 20 s intervals. All data was analyzed with an in-house program written in MATLAB (MathWorks). Data are presented as mean ± SEM. For all LTP experiments, predetermined statistical comparisons were made between the normalized average fEPSP slope 40-60 mins following LTP induction to assess long-lasting changes in synaptic efficacy. Statistical comparisons were performed using two-way ANOVA, and Sidak’s multiple comparisons test was used to compare the same CA region between RGS14 WT and KO animals. Differences between datasets were judged to be significant at p ≤ 0.05. Statistical analyses were performed in GraphPad Prism 7.

To assess whether LTP could be inhibited pharmacologically, slices from RGS14 KO/Amigo2-EGFP+ animals were perfused with either ACSF for controls or ACSF supplemented with either APV (50 µM, Sigma), KN-62 (10 µM, Tocris), or PKI (14-22) amide myristoylated (1 µM, Enzo Life Sciences), to inhibit NMDA receptors, CaMK, or PKA, respectively. For KN-62 experiments, RGS14 KO control slices were perfused with ACSF containing 0.01% DMSO as a vehicle control. Electrophysiological recordings and LTP induction protocol were performed as described above. Data are presented as mean ± SEM. Statistical comparisons were performed using Student’s t test to compare each inhibitor with respective KO control, and differences between datasets were judged to be significant at p ≤ 0.05. Statistical analyses were performed in GraphPad Prism 7.

### Tissue preparation for histology

Adult Amigo2-EGFP mice were anesthetized by isoflurane inhalation and transcardially perfused with 4% paraformaldehyde in PBS. Brains were postfixed for 24h, submerged in 30% sucrose in PBS, and sectioned coronally at 40 µm on a cryostat. Sections were washed in PBS, blocked for at least 1h in 5% normal goat serum (NGS, Vector Labs) diluted in 0.1% PBS-X (0.1% Triton X-100 in PBS) at room temperature, and incubated in primary antibodies diluted in the same buffer overnight. A chicken polyclonal anti-GFP antibody (Abcam) was used at a 1:2,000 dilution to enhance Amigo2-EGFP fluorescence with either a rabbit polyclonal anti-PCP4 antibody (Santa Cruz) or a rabbit polyclonal anti-Wfs1 antibody (ProteinTech). Sections were thoroughly washed in 0.1% PBS-X and incubated in secondary antibodies (Alexa goat anti-chicken 488 and Alexa goat anti-rabbit 568, Invitrogen) diluted at 1:500 for 2h at room temperature. Finally, sections were washed in 0.1% PBS-X and mounted under ProLong Gold Antifade fluorescence media with DAPI (Invitrogen). Sections were then imaged on a Zeiss 710 meta confocal microscope using a 40X oil-immersion lens.

### Organotypic slice preparation

Hippocampal slice cultures were prepared from postnatal day 6-8 RGS14 WT/KO mice as described previously (Stoppini et al., 1991). In brief, hippocampi were dissected and sliced at 320 μm thickness using a tissue chopper. The slices were plated on a membrane filter (Millicell-CM PICMORG50, Millipore). These cultures were maintained at 37 °C in an environment of humidified 95% O_2_ and 5% CO_2_. The culture medium was exchanged with fresh medium every three days. After 7-10 days in culture, neurons were sparsely transfected with ballistic gene transfer (McAllister, 2000) using gold beads (9-11 mg) coated with plasmid containing mEGFP cDNA (10 µg) for sLTP experiments. For RGS14 overexpression experiments, cultured slices from RGS14 KO mice were co-transfected with plasmids containing GFP-tagged RGS14 (20 µg) and CyRFP1 (10 µg, volume marker) cDNAs or mEGFP cDNA (10 µg) as a control. Slices were imaged after 3-12 days following transfection. CA2/CA1 neurons were identified by somatic location within the slice and branching morphology of the apical dendrites.

### Two-photon fluorescence microscopy and two-photon glutamate uncaging

Glutamate uncaging and imaging of live neurons were performed under a custom-built two-photon microscope with two Ti:Sapphire lasers (Chameleon, Coherent). In brief, the lasers were tuned at the wavelength of 920 nm and 720 nm for imaging and uncaging, respectively. The intensity of each laser was independently controlled with electro-optical modulators (Conoptics). The fluorescence was collected with an objective (60x, 1.0 numerical aperture, Olympus), divided with a dichroic mirror (565dcxr) and detected with photoelectron multiplier tubes (PMTs) placed after wavelength filters (ET520/60M-2P for green, ET620/60M-2p for red, Chroma). MNI-caged L-glutamate (4-methoxy-7-nitroindolinyl-caged L-glutamate, Tocris) was uncaged with a train of 4-8 ms laser pulses (2.7-3.0 mW under the objective, 30 times at 0.5 Hz) near a spine of interest. Experiments were performed at room temperature in ACSF solution containing (in mM): 127 NaCl, 2.5 KCl, 25 NaHCO_3_, 1.25 NaH_2_PO_4_, 4 CaCl_2_, 25 glucose, 0.001 tetrodotoxin (Tocris) and 4 MNI-caged L-glutamate, bubbled with 95% O_2_ and 5% CO_2_. We examined secondary/tertiary branches of apical dendrites of CA1 and CA2 pyramidal neurons (located in stratum radiatum) in organotypic cultured hippocampus slices at 10-22 days *in vitro*.

### Calcium Imaging

For Ca^2+^ imaging, we performed whole-cell patch recordings of CA2/CA1 pyramidal neurons in cultured hippocampus slices with the patch pipette containing the Ca^2+^ indicator Fluo-4FF (500 µM; Thermofisher) and Alexa-594 (500 µM) diluted in potassium gluconate internal solution (containing in mM:130 K gluconate, 10 Na phosphocreatine, 4 MgCl_2_, 4 Na_2_ATP, 0.3 MgGTP, L-Ascorbic acid 3, HEPES 10, pH 7.4, 300 mosm). Both dyes were excited simultaneously by a Ti:Sapphire laser (Chameleon, Coherent) at 920 nm. Fluo-4FF (green) and Alexa-594 (red) fluorescence were used to quantify the change in [Ca^2+^] (∆[Ca^2+^]) using the following formula:

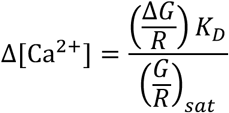

where (G/R)_sat_ is the ratio of green and red fluorescence at a saturating Ca^2+^ concentration measured in the patch pipette (after each day of imaging), and the dissociation constant (K_D_) for Fluo-4FF is 10.4 µM (Yasuda et al., 2004; Lee et al., 2009). Images were acquired using fast-framing two-photon fluorescence microscopy (15.63 Hz frame rate) over a ~4 s baseline before inducing structural plasticity by glutamate uncaging with a train of 4-8 ms laser pulses (2.7-3.0 mW under the objective, 30 times at 0.5 Hz) near a spine of interest. Images were analyzed with MATLAB (MathWorks). Data are presented as the averaged time courses from baseline and the uncaging-triggered average change in [Ca^2+^] (µM) for all 30 glutamate uncaging pulses from baseline ± SEM. To analyze the Ca^2+^ decay kinetics from CA2 spines, the maximum and minimum values from the uncaging-triggered averages were normalized and decay time constants were extrapolated by curve fitting (nonlinear regression) in GraphPad Prism 7.

### Imaging automation

For sLTP experiments images were acquired as a Z stack of five slices with 1 µm separation, averaging five frames/slice. Using multi-position imaging of spines with high-throughput automation, dendritic spines at four positions on separate secondary/tertiary dendrites were imaged simultaneously employing algorithms for autofocusing and drift correction to maintain position and optimal focus during long imaging experiments as recently described (Smirnov et al., 2017). Baseline images were acquired over 5 mins prior to two-photon glutamate uncaging (1 min, 30 pulses at 0.5 Hz) followed by 30 mins of imaging post-uncaging. A 5 min baseline stagger was incorporated to avoid data loss during uncaging events.

### Post hoc immunostaining

Immediately following two-photon imaging experiments, organotypic hippocampal slices were fixed in 4% paraformaldehyde for 30 mins at room temperature. Slices were then washed thoroughly in 0.01M PBS, permeabilized for 15 mins in 0.3% PBS-X (0.3% Triton X-100 in PBS), and thoroughly washed in PBS. Slices were blocked for at least one hour at room temperature in a blocking solution containing 10% NGS (Vector Labs) diluted in 0.1% PBS-X prior to incubation in primary antibodies diluted in the same blocking solution for 42 hours at room temperature. A rabbit polyclonal anti-PCP4 antibody (Santa Cruz) was used at a 1:500 dilution to delineate area CA2 in all slice culture experiments. For sLTP experiments with mEGFP-expressing neurons, a chicken polyclonal anti-GFP antibody (Abcam) was used at a dilution of 1:1,000 to visualize imaged neurons. For Ca^2+^ imaging experiments, Alexa-594 fluorescence was used to identify imaged neurons. Sections were thoroughly washed in 0.1% PBS-X and incubated in secondary antibodies (Alexa goat anti-chicken 488 and Alexa goat anti-rabbit 568, Invitrogen) diluted at 1:500 for 2h at room temperature. After rinsing samples thoroughly in 0.1% PBS-X, slices were optically cleared by incubating in a 60% 2,2′-Thiodiethanol solution (v/v, Sigma) for 30 mins at room temperature (Aoyagi et al., 2015). Intact organotypic slices were imaged in the same clearing solution in glass bottom dishes (Willco) on a Zeiss 880 laser scanning confocal microscope.

### Image and Data Processing

Confocal laser scanning microscope images were processed using FIJI software (NIH v2.0.0). Images were only adjusted for brightness/contrast and cropped for presentation.

### Experimental Design and Statistical Analysis

All statistical analyses were performed in GraphPad Prism 7. Statistical analyses of CA2/CA1 neurons from RGS14 WT/KO mice were made using two-way ANOVA (non-repeated measures) with *post hoc* Sidak’s multiple comparison test between genotypes (i.e. WT CA2-KO CA2 and WT CA1-KO CA1). For pharmacological experiments, LTP in the presence of each inhibitor was compared with paired KO CA2 controls using two-tailed, unpaired t tests. For RGS14 overexpression experiments, CA2 neurons transfected with RGS14-GFP were compared to mEGFP transfected CA2 neurons by unpaired t-test. For RGS14 overexpression experiments in CA1 neurons, all groups were compared by one-way ANOVA followed by Fisher’s LSD test. Complete results of the statistical analyses for each experiment are reported in the results section and sample sizes are noted in the figure legends. In figures * denotes p<0.05, ** denotes p≤0.01, *** denotes p<0.001, **** denotes p<0.0001.

## Results

### Synaptic potentiation can be induced in CA2 in slices from adult RGS14 KO mice

RGS14 has been previously shown to inhibit LTP induction at Schaffer collateral synapses onto CA2 pyramidal neurons by performing whole cell recordings in brain slices prepared from young mice while stimulating in stratum radiatum (Lee et al., 2010). However, the postsynaptic mechanisms underlying RGS14’s actions are poorly understood. Here, we sought to investigate mechanisms by which RGS14 constrains CA2 synaptic potentiation in adulthood when RGS14 levels are highest and stable (Evans et al., 2014). To reliably localize CA2 dendrites in slices, we crossed RGS14 KO mice with an Amigo2-EGFP reporter mouse line that fluorescently labels CA2 pyramidal neurons (Carstens et al., 2016). We validated that this mouse line selectively labels CA2 pyramidal neurons by immunolabeling for the dentate gyrus (DG)- and CA2-enriched protein Purkinje cell protein 4 (PCP4) as well as the CA1 molecular marker Wolframin (WFS1). We found that Amigo2-EGFP fluorescence colocalizes with PCP4 immunoreactivity (Figure 1A) but does not overlap with immunostaining for the CA1 marker WFS1 (Figure 1B).

**Figure 1.**
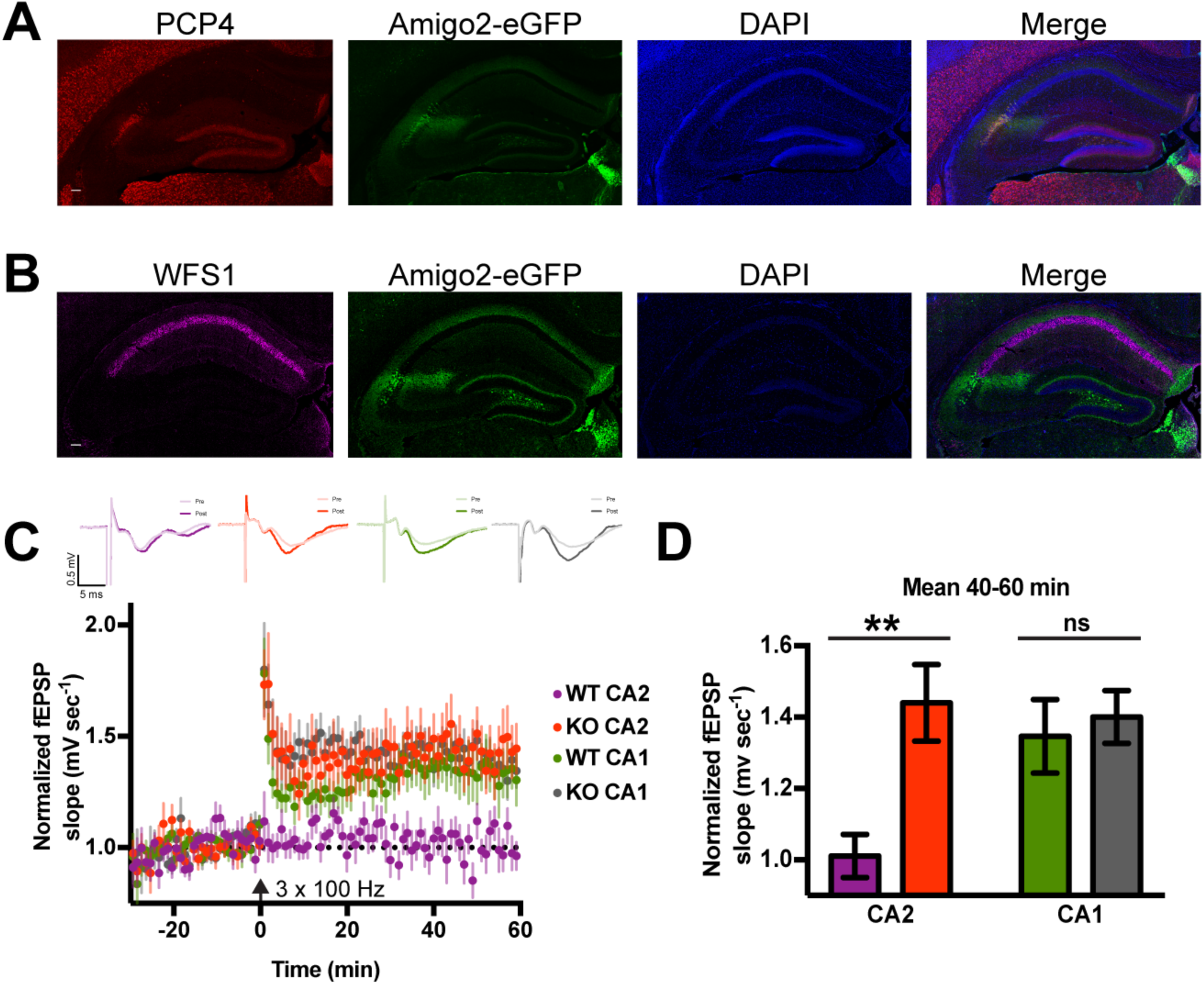
RGS14 KO mice exhibit plasticity of CA2 synapses into adulthood. A. Amigo2-EGFP fluorescence (green) labels CA2 pyramidal neurons as shown by overlap with another CA2 molecular marker PCP4 (red). Scale bar = 100 μm. B. Amigo2-eGFP fluorescence (green) does not colocalize with immunoreactivity for the CA1 pyramidal neuron marker WFS1 (magenta). Scale bar = 100 μm. C. Summary graph of field potential recordings from acute hippocampal slices prepared from adult RGS14 WT and KO mice (both Amigo2-EGFP+) validate that RGS14 KO mice possess a capacity for LTP in CA2 in adulthood (red), which is absent in WT mice (purple). RGS14 WT and KO mice do not differ in CA1 plasticity (green, gray). LTP was induced by high-frequency stimulation (HFS; 3 × 100 Hz) at time 0 (arrow). Data are represented as mean normalized fEPSP slope ± SEM (dotted reference line at 1.0). Sample sizes (in slices/animals) are WT CA2 n = 14/5; KO CA2 n = 16/4; WT CA1 n = 17/4; KO CA1 n = 12/5. Insets (top) are representative traces of field potentials recorded from areas CA2 and CA1 from slices of RGS14 WT and KO mice before (light line) and after (heavy line) LTP induction. D. Quantification of the mean normalized fEPSP slope 40-60 mins following LTP induction (C) with error bars representing SEM. The difference in the degree of LTP induced in RGS14 WT and KO CA2 synapses was significant, whereas no difference was detected between RGS14 WT and KO synapses recorded in CA1. Sidak’s, \*\**p*≤0.01, ns = not significant.

We next performed field potential recordings in acute brain slices prepared from adult RGS14 WT and KO (both Amigo2-EGFP+) mice and replicated previous findings that high-frequency stimulation (3 × 100 Hz) of stratum radiatum induces robust LTP in CA2 neurons of RGS14 KO mice, which is absent in WT mice, and similar to CA1 controls (Lee et al., 2010; Figure 1C). Comparing the mean field excitatory postsynaptic potential (fEPSP) slope averaged from 40-60 minutes after LTP induction, we found that the RGS14 KO CA2 fEPSP slope was significantly larger than WT CA2; LTP induced in CA1 was not significantly different between genotypes (Figure 1D, two-way ANOVA results for RGS14 genotype were F(1,55)=6.64, p=0.0127; results for CA region were F(1,55)=2.478, p=0.1212; and results for interaction were F(1,55)=3.992, p=0.0507. Sidak’s *post hoc* WT CA2–KO CA2 p=0.0036, WT CA1–KO CA1 p=0.9028). Baseline synaptic responses were not altered as input-output curves of fEPSP slope and amplitude were not different in CA2 or CA1 between adult RGS14 WT and KO mice (data not shown).

### CA2 LTP in RGS14 KO mice requires NMDARs, CaMK, and PKA activity

Previous work has shown that LTP in CA2 neurons is naturally suppressed by both robust Ca^2+^ handling mechanisms in spines (Simons et al., 2009) and RGS14 (Lee et al., 2010). To determine if the nascent LTP present in CA2 neurons of RGS14 KO mice requires Ca^2+^ signaling, we next performed the same LTP induction protocol in area CA2 using brain slices prepared from RGS14 KO (Amigo2-EGFP+) mice in the presence of pharmacological inhibitors of Ca^2+^-dependent signaling pathways (Figure 2). The LTP in area CA2 of slices from RGS14 KO mice was effectively blocked by bath application of the NMDAR antagonist APV (50 µM, blue) as well as inhibitors of CaMK (KN-62, 10 µM) or PKA (PKI, 1 µM; Figure 2A-C). Comparing the mean fEPSP slope 40-60 minutes following LTP induction, CA2 plasticity was significantly reduced by each inhibitor relative to paired RGS14 KO controls (Figure 2D; unpaired two-tailed t-tests: for APV t(30) = 2.868, p=0.0075; for KN-62 t(37)=2.59, p=0.0136; for PKI t(37) = 2.502, p=0.0169). These findings indicate that the nascent LTP in CA2 neurons of RGS14 KO mice requires NMDARs and subsequent CaMK and PKA activity, which can be activated – directly or indirectly in the case of PKA – by Ca^2+^ (Figure 2E). Similar mechanisms are thought to support plasticity in CA1 (Malenka et al., 1989; Abel et al., 1997; Otmakhova et al., 2000).

**Figure 2.**
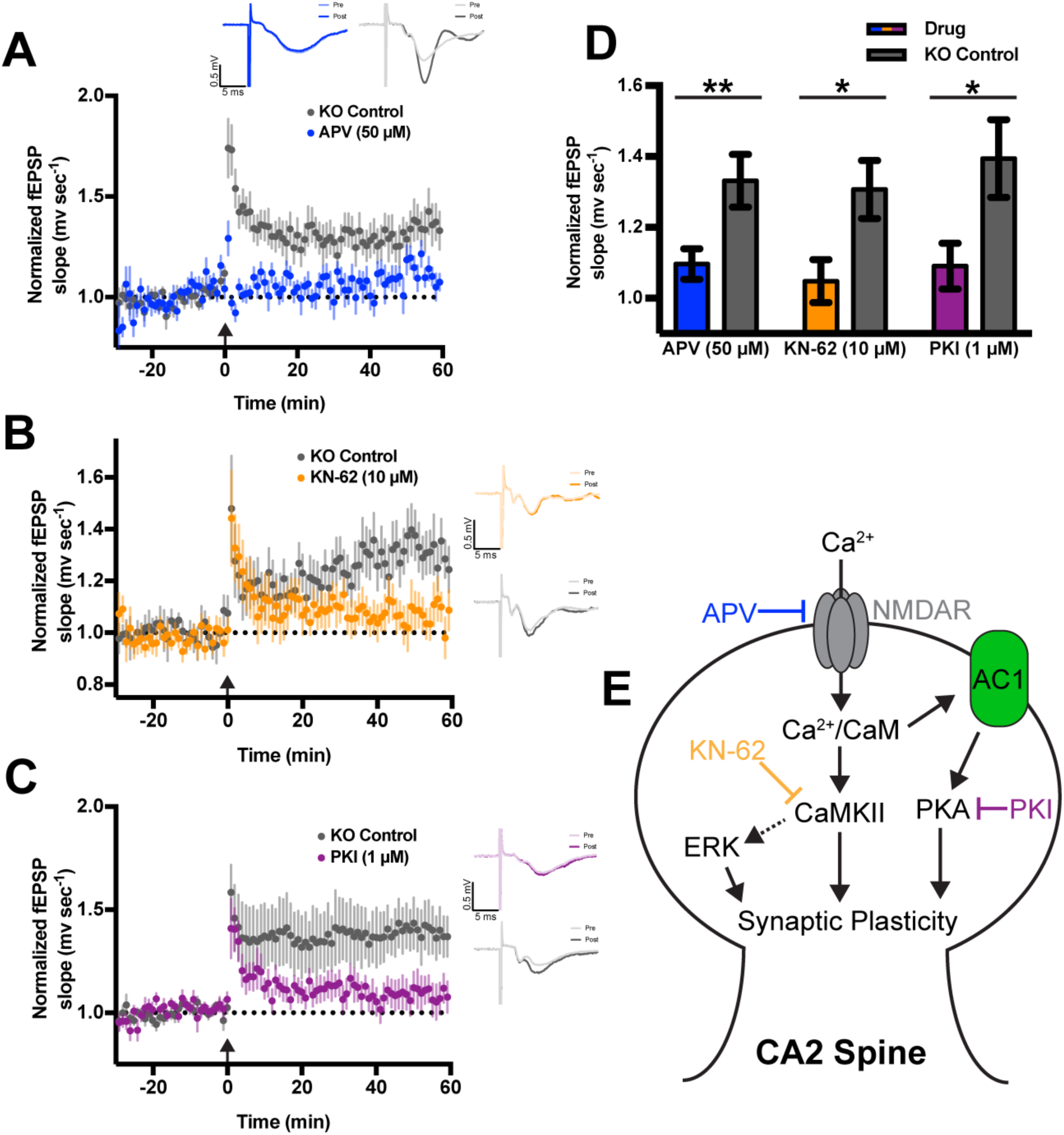
The nascent CA2 LTP in RGS14 KO mice requires Ca^2+^-activated pathways. A. Summary graph of LTP induction experiments performed in area CA2 of RGS14 KO (Amigo2-EGFP+) mice either in the presence (blue) or absence (gray) of bath-applied NMDA receptor antagonist APV (50 μM). LTP was induced by HFS (3 × 100 Hz) in stratum radiatum at time 0 (arrow). Data are represented as mean normalized fEPSP slope ± SEM (dotted reference line at 1.0). Insets (top) are representative traces of field potentials recorded in CA2 stratum radiatum of RGS14 KO mice before (light line) and after (heavy line) LTP induction. B. Summary graph of LTP induction experiments performed in area CA2 of RGS14 KO (Amigo2-EGFP+) mice either in the presence (yellow) or absence (gray) of bath-applied CaMK inhibitor KN-62 (10 μM). C. Summary graph of LTP induction experiments performed in area CA2 of RGS14 KO (Amigo2-EGFP+) mice either in the presence (blue) or absence (gray) of bath applied PKA inhibitor PKI (1 μM). D. Bar graph displaying the mean normalized field potential slope (mV sec^-1^) from data shown in Figure 2A-C, at 40-60 mins following LTP induction with error bars representing SEM. Sample sizes (in slices/animals) are for APV: drug n = 14/10, KO control n = 13/8. For KN-62: drug n = 19/9, KO control n = 16/9. For PKI: drug n = 18/10, KO control n = 16/10. Sample sizes are the same for averaged time courses of each group in Figure 2 A-C. Each inhibitor was compared with paired KO CA2 controls by unpaired two-tailed, t-test; \*\**p* < 0.01, \**p* < 0.05. E. Signaling diagram of a CA2 spine from a RGS14 KO mouse depicting mechanistic targets of the pharmacological inhibitors used in Figure 2A-C.

### RGS14 suppresses structural plasticity of CA2 dendritic spines

After determining that CA2 synapses from RGS14 KO mice utilize similar mechanisms as CA1 neurons to support synaptic potentiation, we next asked if RGS14 might also play a role in activity-dependent spine structural plasticity (sLTP, Figure 3). This enlargement of dendritic spines induced upon synaptic stimulation, is strongly associated with LTP in hippocampus and relies on similar cellular mechanisms (Matsuzaki et al., 2004; Nishiyama and Yasuda, 2015). RGS14 is well positioned to regulate sLTP as it is enriched in CA2 spines and dendrites (Lee et al., 2010); however, no previous studies have examined spine plasticity in CA2 pyramidal neurons. To determine whether RGS14 modulates sLTP in CA2, we cultured hippocampal slices from RGS14 KO mice and WT littermate controls and sparsely transfected neurons with mEGFP using ballistic gene transfer. We then performed two-photon fluorescence microscopy during two-photon glutamate uncaging to induce spine structural plasticity on secondary/tertiary apical dendrites of CA2 and CA1 pyramidal neurons (Figure 3). In order to optimize data collection from each neuron during long experiments, we utilized an automated method to image multiple dendritic spine positions simultaneously (Smirnov et al., 2017). This interface tracks coordinates of multiple dendritic segments on the same neuron and employs autofocus and drift correction algorithms during the experiment to maintain optimal focus. To validate the hippocampal subregion of all neurons imaged in these studies, slices were fixed immediately following two-photon imaging and immunostained for PCP4 to delineate the boundaries of hippocampal area CA2 (Figure 3A, red).

**Figure 3.**
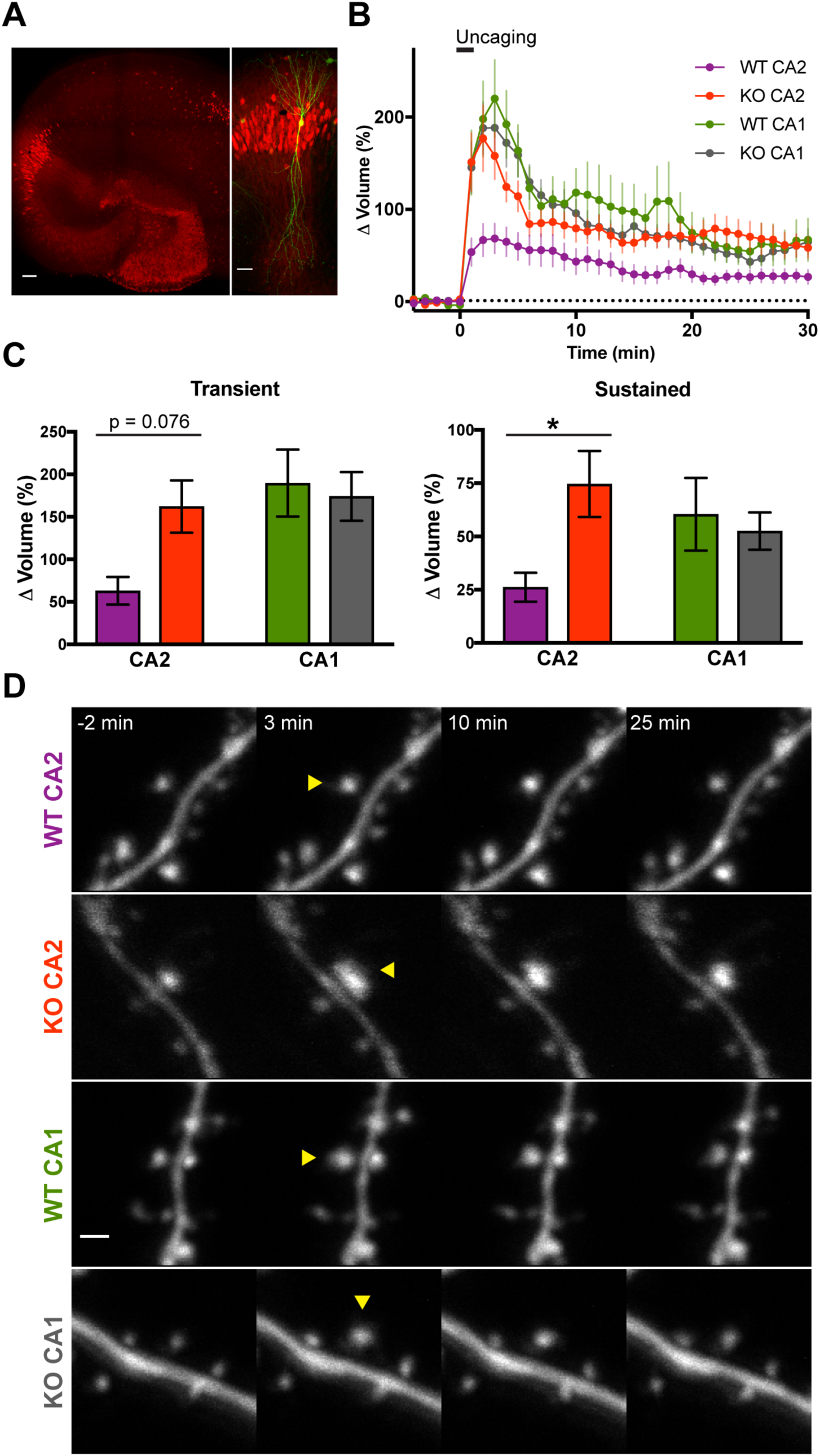
RGS14 suppresses CA2 spine structural plasticity. A. Representative *post hoc* immunostaining to delineate hippocampal region CA2 region after imaging biolistically transfected neurons. *Left*: Organotypic hippocampus slice culture stained for the DG- and CA2-enriched gene PCP4 (red). Scale bar = 100 µm. *Right*: Magnified view of area CA2 in PCP4 immunostained (red) hippocampus on left with a biolistically labeled CA2 pyramidal neuron expressing mEGFP (green). Scale bar = 50 µm. B. Averaged time course of spine volume change during the induction of spine structural plasticity (sLTP) by repetitive two-photon glutamate uncaging (top bar; 30 pulses at 0.5 Hz) in the absence of extracellular Mg^2+^. The number of samples (spines/neurons/animals) for stimulated spines are 19/6/5 for WT CA2, 15/8/6 for KO CA2, 23/7/5 for WT CA1, and 16/6/5 for KO CA1. Sample size applies to Figure 3 B and C. Error bars denote SEM. C. Quantification of the average volume change for stimulated spines during the transient (1-3 mins) and sustained (21-25 mins) phases of sLTP induction. Sidak’s, *p < 0.05. D. Representative two-photon fluorescence images of dendritic spines during sLTP induction in mEGFP-expressing hippocampal pyramidal neurons. Arrowheads indicate location of glutamate uncaging. Scale bar = 1 µm.

Dendritic spines of RGS14 WT CA2 neurons exhibited significantly smaller volume change following repetitive glutamate uncaging to induce sLTP compared to RGS14 KO CA2 neurons or CA1 controls (Figure 3B-D). Comparing the mean spine volume change between samples, the spines of CA2 neurons from RGS14 WT mice trended toward reduced growth in the transient phase of sLTP immediately following uncaging, but the results were not significantly different from the spine volume change of KO CA2 neurons (Figure 3C, left; two-way ANOVA results for genotype were F(1,69) = 1.676, p = 0.1998; results for CA region were F(1,69) = 4.624, p = 0.0350; and results for interaction were F(1,69) = 3.174, p = 0.0792. Sidak’s *post hoc* WT CA2–KO CA2 p=0.0749, WT CA1–KO CA1 p=0.9236.). CA2 spines from slices prepared from RGS14 WT mice exhibited significantly less enlargement compared to RGS14 KO CA2 spines during the later sustained phase of sLTP (Figure 3C, right; two-way ANOVA results for genotype were F(1,69) = 2.172, p = 0.1451; results for CA region were F(1,69) = 0.1954, p = 0.6598; and results for interaction were F(1,69) = 4.198, p = 0.0443. Sidak’s *post hoc* WT CA2– KO CA2 p=0.0360, WT CA1–KO CA1 p=0.8953.). These results indicate that RGS14 naturally restricts the structural plasticity of dendritic spines in CA2 pyramidal cells, similar to the case with functional plasticity.

### RGS14 attenuates spine Ca^2+^ elevations in CA2

Spine Ca^2+^ is critical for the induction of synaptic plasticity (Lynch et al., 1983; Malenka et al., 1988; Harvey et al., 2008), and we found the LTP of CA2 synapses in RGS14 KO mice requires Ca^2+^-driven signaling (Figure 2). Therefore, we investigated if the diminished spine plasticity observed in WT CA2 neurons containing RGS14 could be due to reduced Ca^2+^ levels during glutamate uncaging. Performing two-photon fluorescence microscopy, we monitored Ca^2+^-dependent fluorescence changes in neurons elicited during glutamate uncaging to induce sLTP. Neurons were intracellularly perfused with the synthetic Ca^2+^ indicator Fluo-4FF and the structural dye Alexa-594 to simultaneously monitor Ca^2+^ elevations and spine volume changes elicited by glutamate uncaging. Spine Ca^2+^ responses are presented as averaged time courses (Figure 4 A,B) and the uncaging-triggered average change in Ca^2+^ concentration (∆[Ca^2+^]) across all 30 glutamate uncaging pulses (Figure 4C-E).

**Figure 4.**
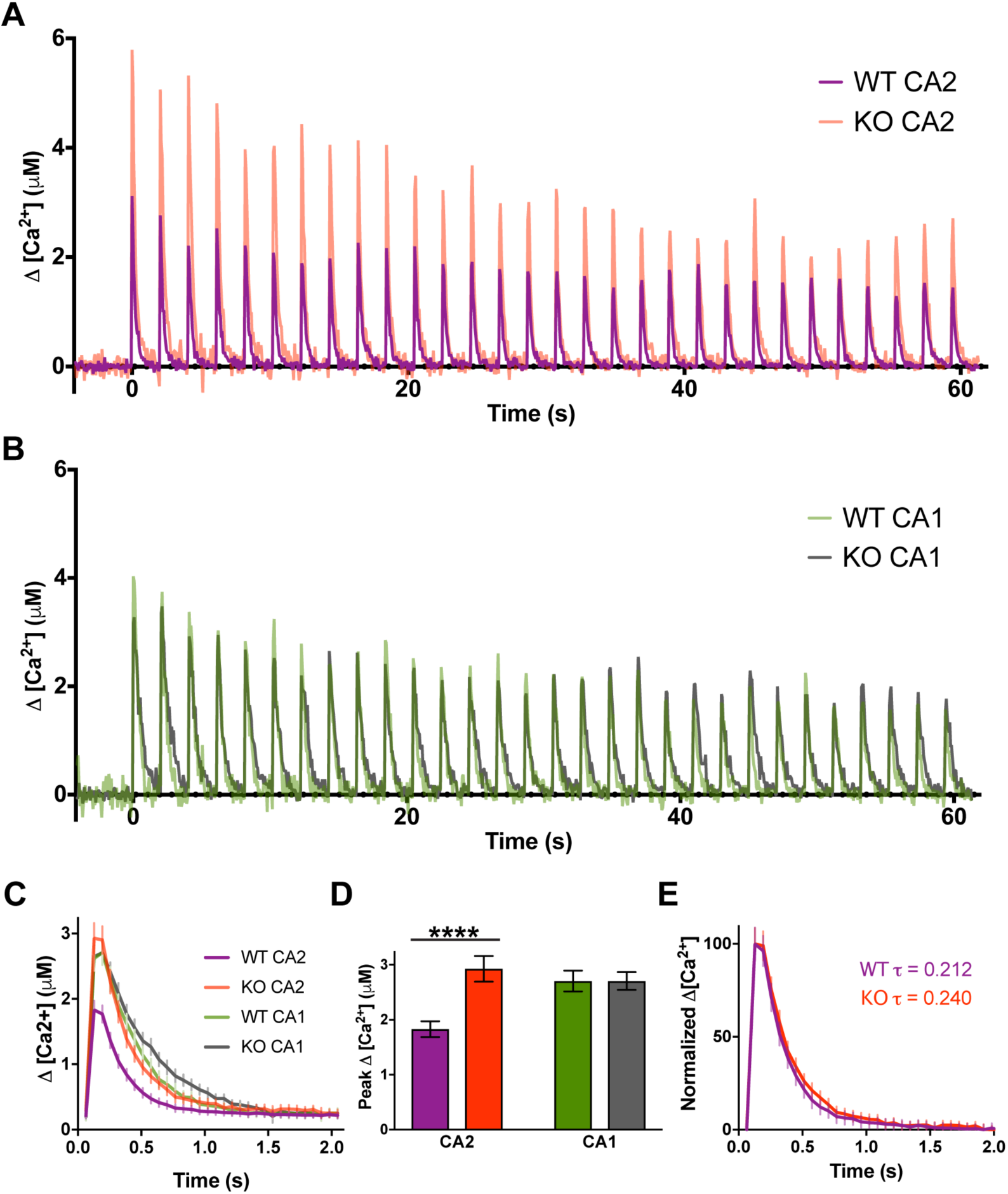
RGS14 restricts Ca^2+^ levels in CA2 spines. A. Averaged time courses of spine Ca^2+^ transients measured with Fluo-4FF during two-photon glutamate uncaging to induce sLTP (30 pulses, 0.5 Hz). Data are shown as the average change in spine Ca^2+^ concentration (∆[Ca^2+^]) from CA2 pyramidal neurons. B. Averaged time courses of spine Ca^2+^ transients measured with Fluo-4FF and Alexa-594 during two-photon glutamate uncaging to induce sLTP (30 pulses, 0.5 Hz). Data are shown as the average change in spine Ca^2+^ (∆[Ca^2+^]) from CA1 pyramidal neurons. C. Uncaging-triggered averages for the change in spine Ca^2+^ concentration during sLTP induction. Error bars denote SEM. D. RGS14 limits CA2 spine Ca^2+^ transients. Bar graphs displaying the average peak ∆[Ca^2+^] ± SEM (****p < 0.0001). The number of samples (spines/neurons/animals) are 40/5/4 for WT CA2, 17/3/3 for KO CA2, 16/3/3 for WT CA1, and 10/2/2 for KO CA1. E. Normalized uncaging-triggered averages for the change in spine Ca^2+^ during sLTP induction reveal similar Ca^2+^ decay kinetics for RGS14 WT and KO CA2 neurons. Error bars denote SEM. WT CA2 average tau is 0.212 s; KO CA2 average tau is 0.240 s.

Uncaging-evoked peak Ca^2+^ elevations were significantly smaller in spines of WT CA2 neurons compared to spines of CA2 neurons from KO littermates, which were comparable to Ca^2+^ responses in CA1 controls (Figure 4A-D; two-way ANOVA results for genotype were F(1,79) = 6.698, p = 0.0115; results for CA region were F(1,79) = 2.367, p = 0.1279; and results for interaction were F(1,79) = 6.675, p = 0.0116. Sidak’s *post hoc* WT CA2–KO CA2 p<0.0001, WT CA1–KO CA1 p>0.9999). Nonlinear regression analysis of normalized uncaging-triggered average Ca^2+^ responses reveals similar decay kinetics in CA2 spines of RGS14 WT and KO mice (Figure 4E; WT CA2 tau = 0.212 s, KO CA2 tau = 0.240 s). Glutamate uncaging also elicited significantly reduced Ca^2+^ transients in WT CA2 dendrites relative to KO CA2 neurons, which were similar to those in CA1 (data not shown). These results demonstrate that spines of CA2 neurons from WT mice display reduced spine Ca^2+^ transients, which correlates with the attenuated spine structural plasticity in the sustained phase, and genetic deletion of RGS14 unmasks synaptic plasticity and restore Ca^2+^ responses to similar levels as those observed in CA2. Together, these data suggest that RGS14 may have a previously unrecognized role in regulating spine Ca^2+^ to constrain CA2 synaptic plasticity.

### Acute RGS14 expression blocks sLTP, and spine plasticity can be recovered by elevating extracellular calcium

To more directly test if there is a causal relationship between RGS14 and the diminished capacity for spine plasticity, we acutely overexpressed GFP-tagged RGS14 (RGS14-GFP) or GFP (control) in organotypic slice cultures prepared from RGS14 KO mice and induced spine plasticity by glutamate uncaging. RGS14-GFP overexpression significantly reduces spine volume enlargement in the sustained phase of sLTP in CA2 pyramidal neurons as expected (Figure 5 A,B; unpaired t-test, ** p = 0.01). RGS14-GFP overexpression similarly blocked transient and sustained phases of spine plasticity in CA1 pyramidal neurons (Figure 5 C,D; Fisher’s LSD test, ** p < 0.01, *** p < 0.001). Given the low probability of transfecting CA2 pyramidal neurons using ballistic gene transfer, we attempted to reverse the plasticity blockade in RGS14-expressing CA1 pyramidal neurons by increasing the extracellular Ca^2+^ concentration from [4 mM] to [8 mM] as a similar manipulation increased spine Ca^2+^ and restored LTP to WT CA2 neurons (Simons et al., 2009). We find that performing sLTP induction in [8mM] external Ca^2+^ significantly rescues the sustained phase of spine plasticity that is otherwise abrogated in RGS14-positive CA1 pyramidal neurons (Figure 5 C,D; Fisher’s LSD test, * p < 0.05, *** p < 0.001). These results demonstrate that RGS14 is able to block long-term structural plasticity in hippocampal neurons, and increasing extracellular [Ca^2+^] restores sustained phase sLTP to RGS14-expressing CA1 neurons. Together, these data suggest that limited Ca^2+^ signaling is central to RGS14’s ability to gate the structural plasticity of dendritic spines as boosting Ca^2+^ can unmask plasticity.

**Figure 5.**
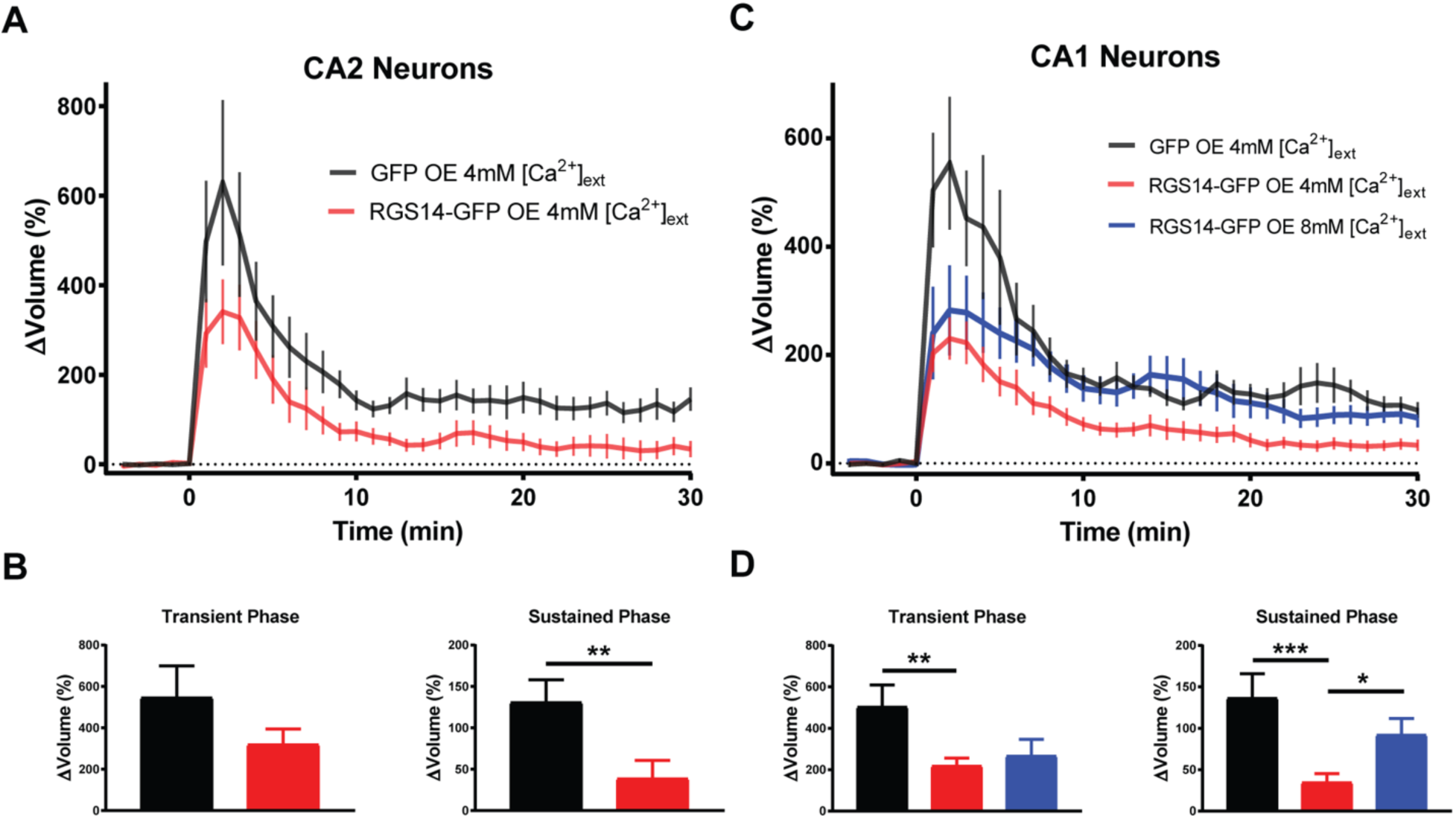
RGS14 expression blocks long-term spine plasticity in CA2 and CA1 neurons lacking RGS14, and high extracellular Ca^2+^ restores structural plasticity to RGS14-expressing neurons. A. Averaged time course of spine volume change in the presence or absence of RGS14 during the induction of spine structural plasticity by repetitive two-photon glutamate uncaging in the absence of extracellular Mg^2+^ in CA2 pyramidal neurons from RGS14 KO mice. The number of samples (spines/neurons/animals) for stimulated spines are 11/4/3 for GFP (black) and 11/4/3 for RGS14-GFP (red). B. Quantification of the average volume change for stimulated spines either in the presence or absence of RGS14 during the transient (1-3 mins) and sustained (21-25 mins) phases of sLTP induction for CA2 neurons. Unpaired t-test, **p = 0.01. C. Averaged time course of spine volume change during the induction of spine structural plasticity in the presence or absence of RGS14 by repetitive two-photon glutamate uncaging in the absence of extracellular Mg^2+^ in CA1 pyramidal neurons from RGS14 KO mice. The number of samples (spines/neurons/animals) for stimulated spines are 11/4/4 for GFP (black) and 23/9/6 for RGS14-GFP 4mM external [Ca^2+^] (red), and 9/4/4 for RGS14-GFP external 8mM [Ca^2+^] (blue). D. Quantification of the average volume change for stimulated spines in the presence or absence of RGS14 during the transient and sustained phases of sLTP induction for CA1 neurons. Fisher’s LSD test, * p < 0.05, **, p < 0.01, *** p < 0.001. Error bars, SEM. Dotted reference line drawn at 0% volume change.

## Discussion

In this study, we have identified a previously unknown role for RGS14 in the regulation of postsynaptic Ca^2+^ signaling in hippocampal CA2 neurons. Specifically, we find that the synaptic potentiation present in CA2 neurons lacking RGS14 requires NMDAR activation along with downstream CaMK and PKA activity, revealing a striking similarity to Ca^2+^-driven mechanisms underlying LTP in CA1 (Malenka et al., 1989; Abel et al., 1997; Otmakhova et al., 2000). We further show that RGS14 naturally inhibits the structural plasticity of CA2 dendritic spines induced by local glutamate uncaging, and loss of RGS14 unleashes robust CA2 spine plasticity observed as long-lasting spine enlargement that is similar to spines in CA1. Although somatic integration of distal and proximal dendritic spines has been examined (Srinivas et al., 2017), to our knowledge this is the first report of the diminished capacity for spine structural plasticity in WT CA2 neurons, similar to the lack of functional synaptic potentiation in stratum radiatum (Zhao et al., 2007). We also demonstrate that the lack of functional and structural plasticity in WT CA2 neurons correlates with attenuated spine Ca^2+^ transients, and plastic spines from CA2 neurons of RGS14 KO mice display larger spine Ca^2+^ transients that are comparable to CA1 controls. Lastly, we find that acute re-expression of RGS14 in brain slices from RGS14 KO mice blocks sustained phase plasticity in CA2 and CA1 pyramidal neurons, and raising the extracellular Ca^2+^ concentration can reverse the blockade of long-lasting sLTP induced by RGS14. Taken together, RGS14 is a critical regulator of CA2 plasticity and Ca^2+^ signaling that seemingly differentiates CA2 pyramidal neurons from the neighboring CA1 subfield, best known for its robust plasticity.

Here, we used a pharmacological approach to reveal the nascent plasticity of CA2 synapses in brain slices of adult RGS14 KO mice requires intact NMDAR, CaMK, and PKA activity (Figure 2), and we previously demonstrated a requirement for ERK activity as well (Lee et al., 2010). These mechanisms are not unique to CA2 pyramidal neurons as they are also thought to support LTP in CA1 as well (Malenka et al., 1989; Abel et al., 1997; Otmakhova et al., 2000). Interestingly, RGS14 engages upstream signaling proteins in each of these kinase cascades: Ca2+/CaM upstream of CaMKs (Evans et al., 2018), Gαi/o subunits that regulate PKA (Cho et al., 2000; Traver et al., 2000; Hollinger et al., 2001), and H-Ras that activates ERK (Kiel et al., 2005; Shu et al., 2010; Vellano et al., 2013). Thus, RGS14 is well positioned to regulate the activities of these pathways, likely in CA2 neurons to naturally gate plasticity therein. Moreover, feedback from these pathways likely regulates the function of RGS14 as it can be phosphorylated by CaMKII (Evans et al., 2018) and PKA (Hollinger et al., 2003). Future studies are necessary to determine how phosphorylation affects RGS14 functions in hippocampal neurons.

Similar to the nascent plasticity of CA2 neurons in RGS14 KO mice shown here and before (Lee et al., 2010), boosting spine Ca^2+^ levels in WT CA2 neurons to levels comparable to those found in CA1 spines, by brief application of high extracellular [Ca^2+^] or blocking Ca^2+^ extrusion through plasma membrane pumps, permits LTP in CA2 pyramidal neurons (Simons et al., 2009). These results demonstrate that CA2 neurons contain the endogenous machinery to support LTP, and led us to hypothesize a common mechanism may mediate the lack of plasticity in CA2. The reduced spine Ca^2+^ transients we observe in WT CA2 neurons is consistent with previous findings that action potential-evoked spine Ca^2+^ transients in CA2 are smaller than in CA3 and CA1, owing to higher rates of Ca^2+^ extrusion and buffering (Simons et al., 2009). These two factors counteract each with regard to decay time, but both reduce peak Ca^2+^ elevation. We found highly similar decay kinetics of spine Ca^2+^ responses between WT and KO CA2 neurons (Figure 4E), which could be attributed to the Ca^2+^ handling that has been previously observed in CA2 spines (Simons et al., 2009). From these data, RGS14 does not likely affect Ca^2+^ extrusion rate or buffering alone, but it may regulate both processes in CA2 spines to reduce peak Ca^2+^ without affecting decay time. One possible mechanism by which RGS14 could regulate spine Ca^2+^ levels is through direct interactions with Ca^2+^/CaM, which has been shown to influence Ca^2+^ dynamics in CA2 and CA1 pyramidal neurons (Simons et al., 2009). Another likely factor contributing to the Ca^2+^ handling properties of CA2 pyramidal neurons, in addition to RGS14, is PCP4/Pep-19, a CA2-enriched protein that modulates calmodulin, which could underlie the robust spine Ca^2+^ extrusion (Simons et al., 2009). Our finding here showing that RGS14 limits peak Ca^2+^ levels in CA2 spines was unexpected, and indicates a shared mechanism of robust Ca^2+^ regulation to inhibit synaptic plasticity in hippocampal CA2.

RGS14 may additionally limit peak Ca^2+^ influx through NMDA receptors, although it has been shown that CA2 NMDAR currents from whole-cell recordings did not differ between RGS14 WT and KO mice (Lee et al., 2010). However, NMDAR Ca^2+^ influx in dendritic spines can be modulated by cAMP-PKA signaling (Skeberdis et al., 2006), which may not be observed by recording current at the soma. NMDAR function and linked signaling in CA2 neurons could be further regulated by the striatal-enriched tyrosine phosphatase (STEP), which is also highly expressed in hippocampal CA2. Future studies are aimed at defining the exact molecular mechanisms by which RGS14 attenuates Ca^2+^ levels in hippocampal CA2 neurons.

Previous studies in hippocampal CA1 neurons have shown that Ca^2+^/CaM inhibitors block both transient and sustained phase of spine plasticity (Matsuzaki et al., 2004) while inhibitors of CaMKII selectively inhibit the sustained phase of sLTP (Matsuzaki et al., 2004; Lee et al., 2009). Therefore, in a WT CA2 spine RGS14 may naturally restrict CaMKII from integrating Ca^2+^ signals – either by regulating the Ca^2+^ levels in spines (Figure 4) or through interactions with Ca^2+^/CaM or CaMKII (Evans et al., 2018) - to achieve full activation necessary to induce long-lasting structural and functional plasticity. Our finding that increasing the concentration of extracellular Ca^2+^ during sLTP induction is sufficient to restore sustained, but not transient, phase plasticity to RGS14-expressing CA1 neurons (Figure 5B) could possibly suggest that this manipulation is sufficient to rescue CaMKII activity levels, but perhaps not those of Ca^2+^/CaM as transient phase plasticity was not restored. Consistently, the LTP of CA2 synapses in brain slices of RGS14 KO mice requires intact CaMKII signaling (Figure 2). Future experiments will investigate if RGS14 regulates the activity of CaMKII in dendritic spines of CA2 neurons.

RGS14 also likely modulates other forms of plasticity in area CA2 in additional to tetanus-induced LTP. RGS14 is well placed to regulate the CA2 synaptic potentiation mediated by oxytocin/vasopressin, which requires postsynaptic Ca^2+^ as well as NMDAR, CaMKII, and ERK activity (Pagani et al., 2015), similar to RGS14 KO mice (Figure 2). RGS14 may also play a role in the caffeine-induced LTP of CA2 synapses, which is G protein- and cAMP-PKA-dependent, but does not require postsynaptic Ca^2+^ (Simons et al., 2012). Lastly, RGS14 could potentially participate in the suppression of CA2 plasticity mediated by perineuronal nets (PNNs, Carstens et al., 2016); acute degradation of PNNs restores plasticity to CA2 in slices, but the relation of RGS14 to PNNs is unknown. While the complex mechanisms that naturally gate LTP in CA2 neurons are not fully understood, it is clear that RGS14 plays a critical role in repressing plasticity therein.

Why would multiple repressive mechanisms exist to block synaptic plasticity in stratum radiatum of area CA2? A study of the expression and localization of RGS14 across development in the postnatal mouse brain showed that RGS14 levels gradually increase with age until reaching highest, stable levels in adulthood (Evans et al., 2014). Given that RGS14 KO mice exhibit markedly enhanced spatial learning and object recognition memory, this developmental upregulation may allow for a period of CA2 plasticity and heightened learning during early postnatal development when spatial navigation, and possibly social bonding with conspecifics, are crucial for survival. We hypothesize that the increase in RGS14 expression after this time allows it to act as a salience filter, such that only specific experiences or forms of memory are encoded as long-term memories by potentiation of CA2 synaptic transmission (Evans et al., 2014).

Although much remains to be explored in the mechanistic underpinnings of restricted plasticity in hippocampal CA2, here we define an unexpected role for RGS14 as a novel regulator of spine Ca^2+^ levels revealing a mechanism upstream of Ras-ERK whereby RGS14 controls synaptic plasticity in CA2. These results provide new insight into the cellular regulation of plasticity in area CA2 and the central role that RGS14 plays in this process.

## Acknowledgements

We thank Jaime Richards for preparing organotypic slice cultures and reagents, David Kloetzer for laboratory management, and Drs. Lesley Colgan and Tal Laviv for assistance with two-photon microscopy.

**Author Contributions:** PE, PP, and DL performed research; PE, PP, and MS analyzed data; PE, SD, JH, and RY designed research; PE wrote and PE, SD, JH and RY edited the paper.

## Notes

**Conflict of Interest:** The authors declare no competing financial interests.

**Funding Sources:** P.R.E. was supported by a predoctoral fellowship from NINDS (1F31NS086174). J.R.H. was supported by NINDS grants 5R01NS37112 and 1R21NS074975. S.M.D. was supported by the Intramural Research Program of NIEHS (Z01ES100221). R.Y. was supported by NIMH grants R01MH080047 and DP1NS096787.

